# Research Process Graph: LLM-Driven Extraction and Hierarchical Organization of Research Logic

**DOI:** 10.64898/2026.06.09.731113

**Authors:** Jing Yang, Manoj Itharajula, Marek Mutwil

## Abstract

Plant biology now publishes thousands of experimental research articles each year, but their core research logic, namely what questions are being asked, with what methods, and what is being found, remains locked inside free text and invisible to systematic analysis. Here we present a structured, 20-year atlas of The Plant Cell in which every paper is converted into a typed, directed Research Process Graph (RPG) of Question (Q), Method (M) and Finding (F) nodes connected by Q→M and M→F edges. A benchmarked large language model pipeline applied to 2,633 Plant Cell research articles published 2005-2026 recovered >110,000 Q/M/F nodes and >126,000 directed Q→M→F chains with>98% precision. A second LLM pass generalises each node into a paper-independent canonical form and assigns it to one of 10 top-level (L1) and ∼90 sub-level (L2) categories for each node type, producing the first comprehensive map of plant-biology research logic at the resolution of individual research questions. The atlas reveals that Plant Cell papers fall into seven canonical paper recipes with characteristic Q→M→F sub-structures, that peripheral experimental techniques have largely turned over while a stable methodological core persisted, and that the strongest correlate of per-PI citation impact is methodological breadth, not productivity or topical breadth. We release the atlas as a public, browsable database with five complementary interfaces: paper views, an LLM-powered research assistant, expert profiles, a taxonomy browser, and a method explorer. The database, available at https://rpg.connectome.tools/, turns the literature into a queryable community resource.

## Introduction

Plant biology has entered an era of genuine multi-scale productivity. The community publishes thousands of experimental research articles per year, each combining genetics, biochemistry, imaging, omics and physiology in increasingly intricate combinations, part of a scientific literature that has expanded near-exponentially for decades (Bornmann and Mutz 2015). The Plant Cell alone has published more than 6,000 research articles since its founding, and the broader plant-biology corpus adds tens of thousands more. As a result, individual researchers can follow only a thin slice of the literature (Landhuis 2016), and the field as a whole lacks any structured, machine-readable view of what is being asked, with what methods, and what is being found across its accumulated work.

The plant community has built powerful curated resources, including TAIR (Lamesch et al. 2012), Araport (Krishnakumar et al. 2015), Phytozome (Goodstein et al. 2012), the Plant Reactome (Fabregat et al. 2018) and plant ontologies (PO, TO and GO)(Ashburner et al. 2001; Cooper et al. 2018),, that organise genes, pathways, traits and organisms. Sequencing repositories (ENA, SRA and GEO)(Leinonen et al. 2011; Clough et al. 2024) and method registries (e.g. protocols.io)(Teytelman et al. 2016) organise data and protocols. But none of these capture what a paper actually did: the research (Q)uestion that motivated each experiment, the (M)ethod chosen to address it, and the (F)inding it produced. This experimental logic, the directed Q → M → F chain that defines each piece of work, is currently locked inside free-text articles and unavailable for systematic analysis.

Large language models (LLMs) have matured to the point where they can read scientific articles and produce structured, typed output with high precision (Lim et al. 2025). Recent applications have extracted gene-function descriptions (Hu et al. 2025), materials data and synthesis methods (Dagdelen et al. 2024; Polak and Morgan 2024), drug-target pairs (Luo et al. 2022) and disease-phenotype links (Yang et al., 2024), and entity-relation mining now spans tens of millions of articles (Wei et al., 2024). Plant biology has begun to adopt the same tools: PlantConnectome used large language models to mine over 71,000 plant articles into a knowledge graph of nearly 5 million functional relationships between genes, metabolites, tissues and other entities (Lim et al., 2025), retrieval-augmented agents such as PlantGPT now answer functional-genomics questions grounded in the literature (Zhang et al., 2025), and the promise of combining language models with knowledge graphs to accelerate gene-function discovery has been set out in recent reviews (Lam et al., 2024; Sunil et al., 2024). A parallel effort uses LLM agents to conduct research end-to-end: formulating hypotheses, running experiments and drafting the paper as in The AI Scientist (Lu et al., 2024) and AI Scientist-v2 (Yamada et al., 2025). Yet all of these efforts capture entities and pairwise relations rather than the directed research logic of a paper.

Here, we use LLMs to extract a more demanding object: the Research Process Graph (RPG) for each paper, composed of typed Question (Q), Method (M), and Finding (F) nodes connected by directed Q→M and M→F edges. Each RPG is a small, machine-readable map of the experimental reasoning of one paper. We applied this pipeline to 2,633 Plant Cell research articles spanning 2005–2026, recovering 110,295 typed Q/M/F nodes and 126,388 directed Q→M→F chains at >98% precision against a manually annotated gold standard. A second LLM pass then generalised and organised every node into a hierarchical Q/M/F taxonomy. The resulting atlas, to our knowledge the first structured map of research logic, lets us characterise the field on its own terms: which types of questions are asked, how they couple to the methods used and the findings produced, how papers fall into a small set of recurring research “recipes”, and how a principal investigator’s methodological breadth relates to research impact.

We release the atlas as a public, browsable database with complementary interfaces for exploring papers, researchers, techniques and the taxonomy, including an LLM-powered assistant that returns literature-grounded research prospectuses for any natural-language question. Because the RPG framework is journal- and field-agnostic, The Plant Cell serves here as a proof of concept that an entire branch of biology can now be mapped at the resolution of its individual research questions.

## Results

### An LLM pipeline extracts Question/Method/Finding knowledge graphs from 2,633 Plant Cell papers with >98% precision

To capture the experimental logic of a paper in a structured form, we designed a pipeline that represents each article as a knowledge graph of Research Questions (Q), Methods (M) and Findings (F) connected by directed Q→M and M→F edges (Figure 1A). Full-text articles were filtered to retain experimental research papers (Table S1), then processed with GPT-5 via the OpenAI Batch API using a structured extraction prompt, producing one JSON record of typed nodes and edges per paper. A representative Q→M→F triple from a rice leaf-angle study illustrates the output (Figure 1A, inset).

**Figure 1.**
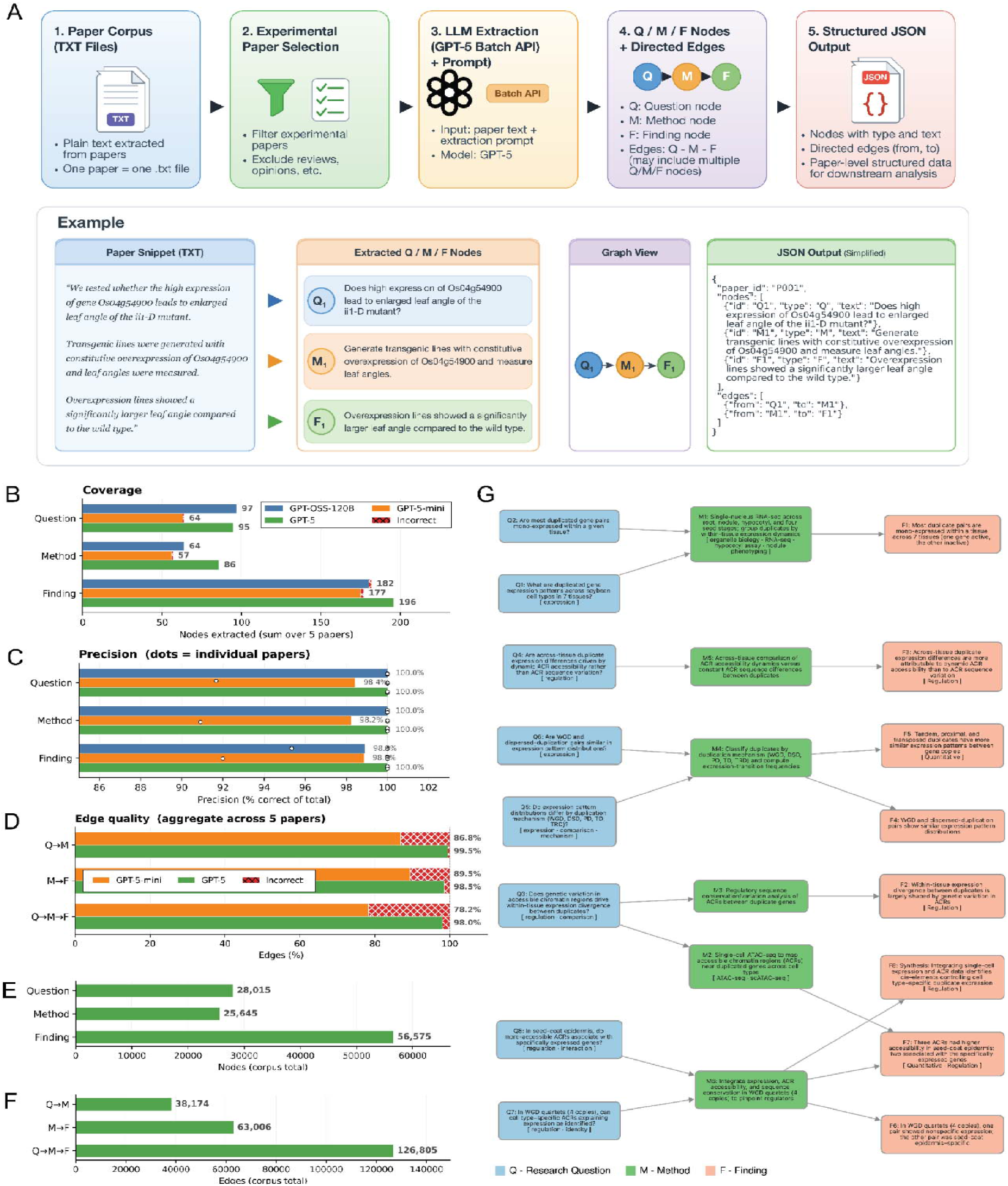
Q/M/F extraction from experimental plant biology papers. (A) Overview of the extraction pipeline. Plain-text paper files (step 1) are filtered to keep experimental research articles only (step 2). Each paper is sent through GPT-5 via the OpenAI Batch API together with an extraction prompt (step 3) that asks the model to identify the paper’s Research Questions (Q), Methods (M) and Findings (F) and the directed edges between them (step 4). The output is a structured JSON record per paper containing typed nodes, their original text, and Q→M and M→F edges (step 5). The boxed example illustrates a single Q→M→F chain drawn from one rice leaf-angle study. (B) Node coverage: number of Q, M and F nodes extracted by GPT-OSS-120B, GPT-5-mini and GPT-5, summed across the five benchmark papers. The red hatched fraction shows incorrect nodes. (C) Node-level precision: percent of extracted nodes judged correct by manual review; dots indicate per-paper precision. (D) Edge quality across the same five papers for GPT-5-mini and GPT-5: precision of Q→M edges, M→F edges, and full Q→M→F chains. (E, F) Corpus statistics across the full set of 2,633 Plant Cell papers extracted with GPT-5. (E) Total node counts by type. (F) Total edge. (G) Example knowledge graph from a representative paper on gene duplicate expression. Q nodes (blue) are research questions, M nodes (green) are the experimental methods used to address them, and F nodes (orange) are the findings supported by those methods. Each method node carries a short list of recognised technique tags (in square brackets), and each F node a category tag. Arrows indicate the directed Q→M and M→F edges extracted by the model.

Before applying the pipeline at scale, we benchmarked its accuracy against five manually annotated papers and compared three OpenAI models that span a range of capability and cost: the frontier GPT-5, the lower-cost GPT-5-mini and the open-source GPT-OSS-120B. All three were evaluated under an identical prompt and structured-output interface, so that any differences in performance and costs can be attributed to the models themselves rather than to variation in prompting or implementation. All three models extracted most nodes correctly in terms of both number of nodes and edges and precision (Table S2-11), but GPT-5 performed best, reaching approximately 100% precision for Q, M and F nodes while also recovering the most nodes overall (95 Q, 86 M and 196 F across the five papers; Figure 1, B and C)(Table S2-6). The advantage of GPT-5 was clearest at the level of edges (Table S7-11), where it produced at least 98% correct Q→M, M→F and full Q→M→F chains, whereas GPT-5-mini fell to 78% correct for full chains (Figure 1D). We therefore selected GPT-5 for corpus-wide extraction.

Applying the GPT-5 pipeline to the full set of 2,633 Plant Cell papers yielded 110,235 nodes (28,015 Q, 25,645 M and 56,575 F). Papers contained, on average, approximately 10 Q and 10 M nodes, but roughly twice as many F nodes, confirming that Findings are the most numerous node type and that each question or method typically supports multiple findings (Figure 1E). The same asymmetry was evident among edges: M→F edges (63,006) outnumbered Q→M edges (38,174), indicating that a single method generally supports more findings than a single question recruits methods (Figure 1F). Composing the two edge types produced 126,805 complete Q→M→F chains, which serve as the basic units for all downstream analyses.

The resulting per-paper graphs capture the non-linear structure of real studies. In an example RPG from a study of duplicated-gene expression in soybean (Fattorini 2026), some questions are addressed by multiple methods, a single method is shared across several questions, and individual findings integrate evidence from more than one method (Figure 1G).

### Generalising and clustering Q/M/F nodes produces a hierarchical taxonomy of plant biology research

The per-paper graphs use the specific language of each study, which makes the corpus difficult to analyse as a whole. To uncover the categories of questions, methods and findings recurring across papers, we developed a second pipeline that converts each Q/M/F text into a generalised form: a paraphrase that removes paper-specific entities such as gene names, organisms and experimental scales while preserving the underlying scientific intent (Figure 2A). Because this task is simpler than constructing the RPG itself, we used the less expensive GPT-5-mini with a normalisation prompt, supplying each paper’s extracted list of abbreviations as side input so that acronyms such as cryo-EM and MET1 were resolved before generalization. Each record retains the original text, its generalised form and the modifications applied.

**Figure 2.**
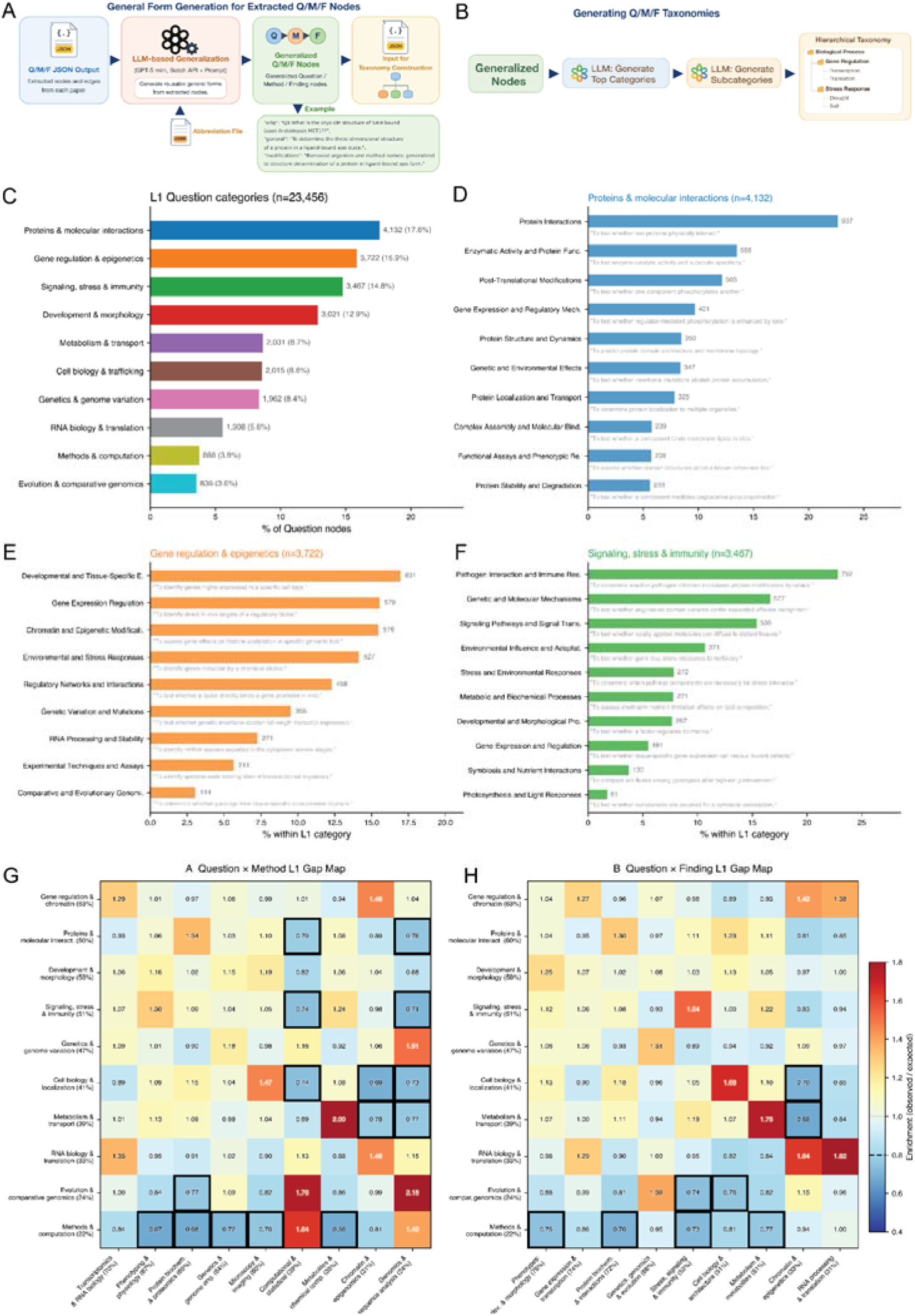
Generalisation and hierarchical taxonomy of Q/M/F nodes across the Plant Cell corpus. (A) Generalisation pipeline. Each paper’s raw Q/M/F nodes are passed through GPT-5-mini with a normalisation prompt that strips paper-specific entities while preserving scientific intent. A per-paper abbreviation file (extracted from the original text) is supplied as side input so acronyms are resolved before generalisation. The output JSON keeps the original text (orig), the generalised form (general) and a note of the modifications applied. (B) Two-stage taxonomy construction. Unique generalised nodes of each type are fed to an LLM that proposes top-level categories (L1); a second LLM call within each L1 proposes sub-categories (L2). Generalised nodes are then assigned to one L1 × L2 cell. (C) L1 distribution of Question nodes (n = 23,456). (D–F) L2 sub-structure within the three largest Q-L1 categories, showing the number of Q nodes per L2 and one representative generalised question (italic): (D) Proteins & molecular interactions (n = 4,132), (E) Gene regulation & epigenetics (n = 3,722), (F) Signaling, stress & immunity (n = 3,467). (G, H) L1 coupling matrices, computed at the paper level. Rows are Q-L1 categories; columns are matched M-L1 categories in (G) and F-L1 categories in (H). Cells show the enrichment ratio (observed/expected under independence); black-outlined cells indicate statistically over- or under-represented pairings. Row labels include the marginal share of papers touching each Q-L1.

To organise the generalized nodes per type into a navigable structure, we applied a two-stage LLM categorisation pipeline (Figure 2B). An LLM first proposed top-level (L1) categories from the generalised nodes, and then, in a second call, proposed sub-categories (L2) within each L1 category. Every generalised node was then assigned to a single L1 by L2 cell, and the procedure was iterated until the assignments stabilised.

Across the 23,456 Question nodes, this procedure produced 10 L1 categories that summarise what the field asks (Figure 2C, the L1 and L2 categories per paper are found in Table S1). Proteins and molecular interactions was the most studied category (17.6%), followed by Gene regulation and epigenetics (15.9%), Signaling, stress and immunity (14.8%) and Development and morphology (12.9%); together, the long tail of methods, computation and evolution questions accounted for less than 8% of the corpus. The L2 sub-structure of the largest categories recapitulated familiar research programmes. Within Proteins and molecular interactions (n = 4,132), protein-protein interaction questions dominated (937 nodes), followed by enzymatic activity and protein function, post-translational modifications, protein structure and dynamics, and protein localisation, capturing the classic biochemistry questions (Figure 2D). Gene regulation and epigenetics questions (n = 3,722) were led by developmental and tissue-specific expression, gene expression regulation, and chromatin and epigenetic modifications, with environmental and stress responses and regulatory-network questions making up the remainder (Figure 2E). Within Signaling, stress and immunity (n = 3,467), pathogen interaction and immune responses formed by far the largest L2 (792 nodes), identifying plant pathology as a defining sub-field (Figure 2F). Thus, this analysis reveals the major categories and sub-categories for Q (Figure S1), M (Figure S2) and F (Figure S3) nodes.

With questions, methods and findings placed in a shared taxonomy, we could ask how they combine within papers. Comparing Question-L1 and Method-L1 categories co-occurring in the same paper by enrichment analysis (observed over expected co-occurrence under independence) revealed strong, expected coupling along the diagonal: Metabolism and transport questions paired with metabolite and chemistry methods (ratio 2.00), Evolution and comparative genomics questions with genomics and sequence methods (2.18), and Cell biology and localisation questions with microscopy and imaging methods (1.47)(Figure 2G). The depleted cells were equally informative, marking method-question incompatibilities: Signaling, stress and immunity questions were systematically under-coupled to genomics and sequence methods (0.71); Cell biology and localisation questions avoided genomics and sequence methods (0.73) and chromatin methods (0.69); and Metabolism and transport questions showed the most asymmetric profile, strongly enriched for metabolite methods but depleted for chromatin and epigenomics (0.77).

The same analysis applied to Question-L1 and Finding-L1 categories showed that questions generally produce findings in the same domain (Metabolism and transport questions to Metabolism findings, ratio 1.75; RNA biology and translation to RNA processing, 1.82; Signaling, stress and immunity to Stress and signalling, 1.54), alongside some productive cross-talk, such as RNA biology & translation questions and Chromatin and epigenetics findings (1.64) (Figure 2H). Conversely, certain domain boundaries were rarely crossed: Cell biology and localisation questions almost never produced Chromatin and epigenetics findings (0.70); Methods and computation questions were decoupled from most biology-flavoured findings (ratios 0.70 to 0.86); and Proteins and molecular interactions questions were notably depleted in Chromatin and epigenetics findings (0.81). Taken together, the enriched diagonal and the depleted off-diagonal define the implicit experimental boundaries of the field, mapping which question types are addressed by which methods and which findings are unlikely to follow.

### Seven research recipes structure the Plant Cell corpus and predict per-PI impact

If questions, methods and findings couple in predictable ways, papers as a whole should occupy a limited region of the possible space. To test this, we represented each paper as a 28-dimensional L1 composition vector of block-normalised Q-L1, M-L1, and F-L1 shares, clustered the 2,558 papers using k-means (k = 7), and projected the results with UMAP (Figure 3A). The papers resolved into seven well-separated clusters, which we term paper recipes, each defined by a dominant Q-L1 to M-L1 to F-L1 triad: gene regulation through transcriptomics to gene expression (R1), protein interactions through protein biochemistry to protein biochemistry (R2), development through microscopy to developmental phenotypes (R3), signaling and stress through phenotyping to stress signalling (R4), metabolism through metabolites to metabolism (R5), genetics through computational methods to genetics and evolution (R6), and cell biology through microscopy to cell biology (R7).

**Figure 3.**
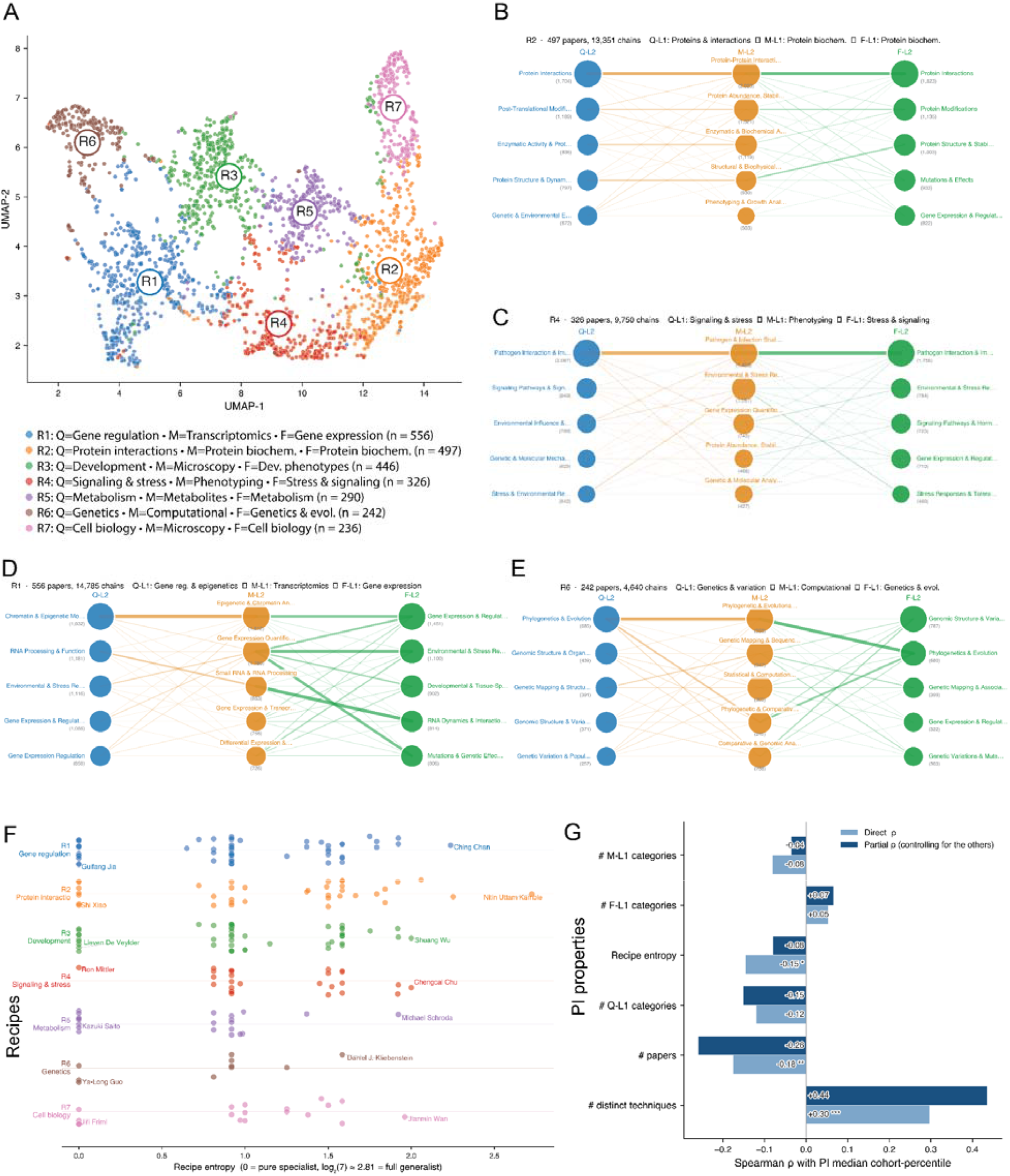
Paper recipes, PI specialisation and predictors of impact. (A) UMAP projection of 2,558 Plant Cell papers, each represented as a 28-dimensional L1 composition vector (block-normalised Q-L1, M-L1, F-L1 shares). Colour and R-label mark the k-means cluster (k = 7); legend gives the dominant Q-L1 → M-L1 → F-L1 triad and the cluster size. (B–E) Chain networks for four representative recipes: (B) R2, (C) R4, (D) R1, (E) R6, built from real Q→M and M→F edges in the paper JSONs and projected onto L2 sub-categories. Nodes are the top 5 Q-L2 (blue), M-L2 (orange) and F-L2 (green) within the recipe; node size is proportional to the number of chains touching that L2; edge thickness is proportional to the chain count between connected L2 pairs. Panel headers list the recipe ID, paper count, total chain count, and the defining Q-L1 → M-L1 → F-L1 triad. (F) PI specialisation by dominant recipe (n = 218 PIs with ≥3 corpus papers). Each dot is one PI, placed along the x-axis by the Shannon entropy of their recipe distribution (0 = pure specialist, log_2_(7) ≈ 2.81 = full generalist). PIs are stacked in seven horizontal bands by their dominant recipe; the most extreme specialist and most generalist of each band are labelled. (G) Spearman correlations between PI median cohort-percentile citation score and six candidate predictors. Light bars: direct ρ; dark bars: partial ρ (each predictor controlling for all five others). Asterisks on the direct ρ indicate significance (* p < 0.05, ** p < 0.01, *** p < 0.001).

To resolve the experimental substructure within each recipe, we mapped the real Q→M→F edges from the per-paper records to their L2 subcategories. R2 (Proteins; 497 papers, 13,351 chains) was anchored by protein-interaction questions answered with protein-protein interaction and abundance or stability assays, yielding findings dominated by protein interactions and modifications (Figure 3B). R4 (Signalling and stress; 326 papers, 9,750 chains) showed a strong pathogen-interaction spine, in which pathogen-related questions were answered by pathogen and infection assays, running alongside a narrower environmental-stress branch (Figure 3C). R1 (Gene regulation; 556 papers, 14,785 chains) contained two distinct sub-flavours within a single recipe: a chromatin and epigenetics branch, in which chromatin and epigenetic modification questions were addressed by epigenetic and chromatin analyses, and an RNA-biology branch, in which RNA-processing questions were addressed by small-RNA and RNA-processing methods (Figure 3D). R6 (Genetics and comparative genomics; 242 papers, 4,640 chains) consistently routed questions in phylogenetics and evolution through phylogenetic and comparative methods to produce genomic variation findings, recapitulating the canonical comparative genomics workflow (Figure 3E, Figure S4 for all chain networks).

We next asked how individual researchers distribute their work across these recipes. Computing the Shannon entropy of each principal investigator’s recipe distribution for the 218 PIs with at least three corpus papers, and stratifying by dominant recipe, showed that every recipe contained both pure specialists (entropy = 0, for example Kazuki Saito in R5 and Jiří Friml in R7) and broad generalists (entropy > 2, for example Nitin Uttam Kamble in R2 and Chengcai Chu in R4)(Table S12); neither extreme was absent from any recipe (Figure 3F).

Finally, we asked which PI-level features predict research impact, as measured by citations their papers accumulated. Correlating each PI’s median cohort-percentile citation score with six candidate predictors revealed that the number of distinct techniques a PI used was the strongest positive predictor (direct Spearman ρ = +0.30, partial ρ = +0.44), whereas the number of papers was significantly negative (partial ρ = −0.26), and recipe entropy had almost no unique effect once the number of papers and number of techniques were controlled (partial ρ = −0.08) (Figure 3G). Thus, methodological breadth, rather than productivity or topical breadth, is therefore the feature most closely associated with per-paper impact in this corpus.

### Method composition turns over, but methodological breadth per paper stays constant

The recipes capture the methodological identity of papers, but not how that identity has changed over time. To identify which techniques have entered or left plant biology, we extracted the methodological vocabulary of every paper from its M-node text using a curated dictionary of approximately 305 techniques (Table S1), and compared the share of papers using each technique in an early window (2005 to 2010) and a recent window (2020 to 2024) (Figure 4A). The shifts were pronounced. Several techniques rose sharply: RNA-seq increased from 0.3% to 34% of papers, CRISPR from 0 to 21%, and ChIP-seq, IP-MS, GWAS, split-luciferase, BiFC, Y1H/Y2H and small-RNA-seq each rose at least two-fold. Others declined just as steeply: PCR fell from 70% to 24%, RT-PCR from 49% to 7%, Northern blot from 21% to 2%, microarray from 18% to 1%, with T-DNA and GUS reporter assays also dropping sharply. A core set of techniques, however, remained in use throughout, each appearing in more than 25% of papers across the full 20 years: Western blot, RT-qPCR, GFP and fluorescent tagging, confocal microscopy and overexpression.

**Figure 4.**
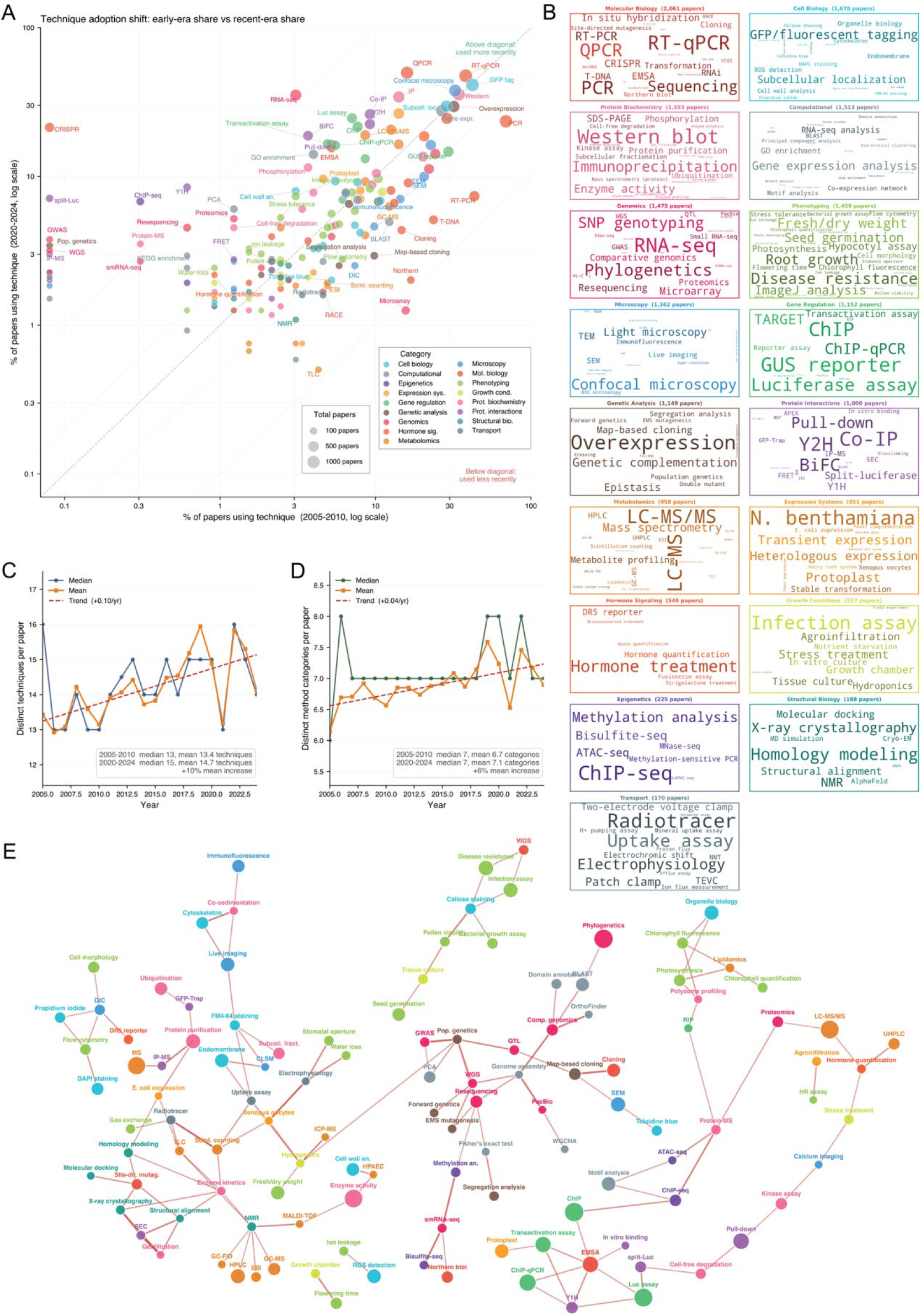
Twenty years of methodological change in Plant Cell papers. (A) Technique adoption shift, 2005-2010 vs 2020-2024. Each dot represents one technique (≥30 corpus papers); x and y show the share of papers using the technique in each window, on a log scale. Point size is proportional to the total number of papers using the technique; colour indicates the method category. The diagonal indicates no change; the two parallel dotted lines indicate 2- and 10-fold changes. (B) Per-category technique word-clouds for the 17 method categories. Within each box, font size is proportional to the number of corpus papers that use the technique; the category-level paper count is shown in the box header. (C) Distinct techniques per paper, 2005-2024. Lines show the per-year median (blue), per-year mean (orange), and a linear trend fit to the mean (red, dashed). Inset gives the median, mean and percent change between the two reference windows. (D) Distinct method categories per paper (of 17 possible), 2005-2024, with the same conventions as in (C). (E) Technique co-occurrence network. Nodes are techniques used in ≥30 papers; edges connect technique pairs whose paper-level co-occurrence enrichment (log_2_(observed/expected)) exceeds a chosen threshold. Node size is proportional to the number of corpus papers using the technique; node colour denotes the method category.

To summarise the methodological signature of each of the 17 method categories, we generated per-category word clouds in which the size of each technique is proportional to the number of papers using it (Figure 4B). The largest category, Molecular Biology (2,061 papers), was dominated by qPCR, RT-qPCR, PCR and CRISPR; Cell Biology by GFP tagging and subcellular localisation; Microscopy by confocal microscopy; Metabolomics by LC-MS, LC-MS/MS and metabolite profiling; and Gene Regulation by GUS reporter, luciferase assay, ChIP and transactivation assays, giving each category a recognisable identity.

We next asked whether this turnover was accompanied by an increase in methodological complexity. Counting the distinct techniques per paper across 2005 to 2024 revealed only a modest increase, from a median of 13 in 2005 to 2010 to 15 in 2020 to 2024 (mean 13.4 to 14.7, a 10% rise, trend +0.10 techniques per year), indicating that Plant Cell papers have always been methodologically dense and have broadened only slightly (Figure 4C). The same was true at the level of method categories: papers consistently spanned roughly 7 of the 17 categories with little change over two decades (median 7 throughout; mean 6.7 to 7.1, a 6% rise, trend +0.04 categories per year) (Figure 4D). Taken together, these panels show that the defining trend of the past 20 years in plant science has been compositional turnover rather than expansion: each generation of papers replaced old techniques with new ones at nearly a one-to-one rate, while the core toolkit and the overall breadth of methods per paper remained largely stable.

Finally, we asked how individual techniques relate to one another. We built a technique co-occurrence network in which nodes are techniques used in at least 30 papers, and edges connect techniques that appear together more often than expected by chance. Edges are weighted by the paper-level log2 enrichment of observed over expected co-occurrence, with nodes coloured by method category (Figure 4E). Several biologically coherent modules emerged from the data alone: a biochemistry and enzymology core (NMR, X-ray crystallography, molecular docking, size-exclusion chromatography and enzyme kinetics), a plant pathology and phenotyping cluster (disease resistance, infection assays, bacterial growth and hypersensitive-response assays), a cell-biology imaging hub (live imaging, FM4-64, confocal microscopy, immunofluorescence and cytoskeleton imaging), a comparative-genomics cluster (GWAS, whole-genome sequencing, PacBio, OrthoFinder and phylogenetics) and a gene-regulation cluster (ChIP, EMSA, transactivation assays, luciferase, Y1H, ChIP-seq and bisulfite-seq). The techniques bridging these modules, such as mass spectrometry linking biochemistry to proteomics and scanning electron microscopy linking imaging to phenotyping, identify the methods that span multiple research traditions.

### A public RPG database exposes the corpus through five complementary interfaces: paper views, an AI research assistant, expert profiles, taxonomy browser and method explorer

To make our findings usable by the community, we built a public, browsable database that exposes the RPG data through five complementary interfaces. To make the project concepts more easily accessible to the public and their AI agents, we developed an Obsidian wiki containing the project description at https://publish.obsidian.md/mutwillab/Homepage/Publication+Vaults/RPG/README.

#### Paper panel

The paper view presents each article as a structured record, tabulating its extracted Research Questions, Methods, and Findings in three colour-coded columns, illustrated here for a wheat study of the *Puccinia striiformis* races CYR23 and CYR31 (Figure 5A)(Ma et al. 2023). The same nodes are also rendered as a compact Q→M→F graph, making the paper’s internal logic explicit and showing which method addresses which question and which finding it supports (Figure 5B). To help users locate functionally related work, the database computes paper-to-paper similarity based on shared Q-L2, M-L2, and F-L2 tags; users can reweight the Q, M, and F contributions and the specificity of the L2 tags to bias the ranking and retrieve related papers along with their overlap profile (Figure 5C).

**Figure 5.**
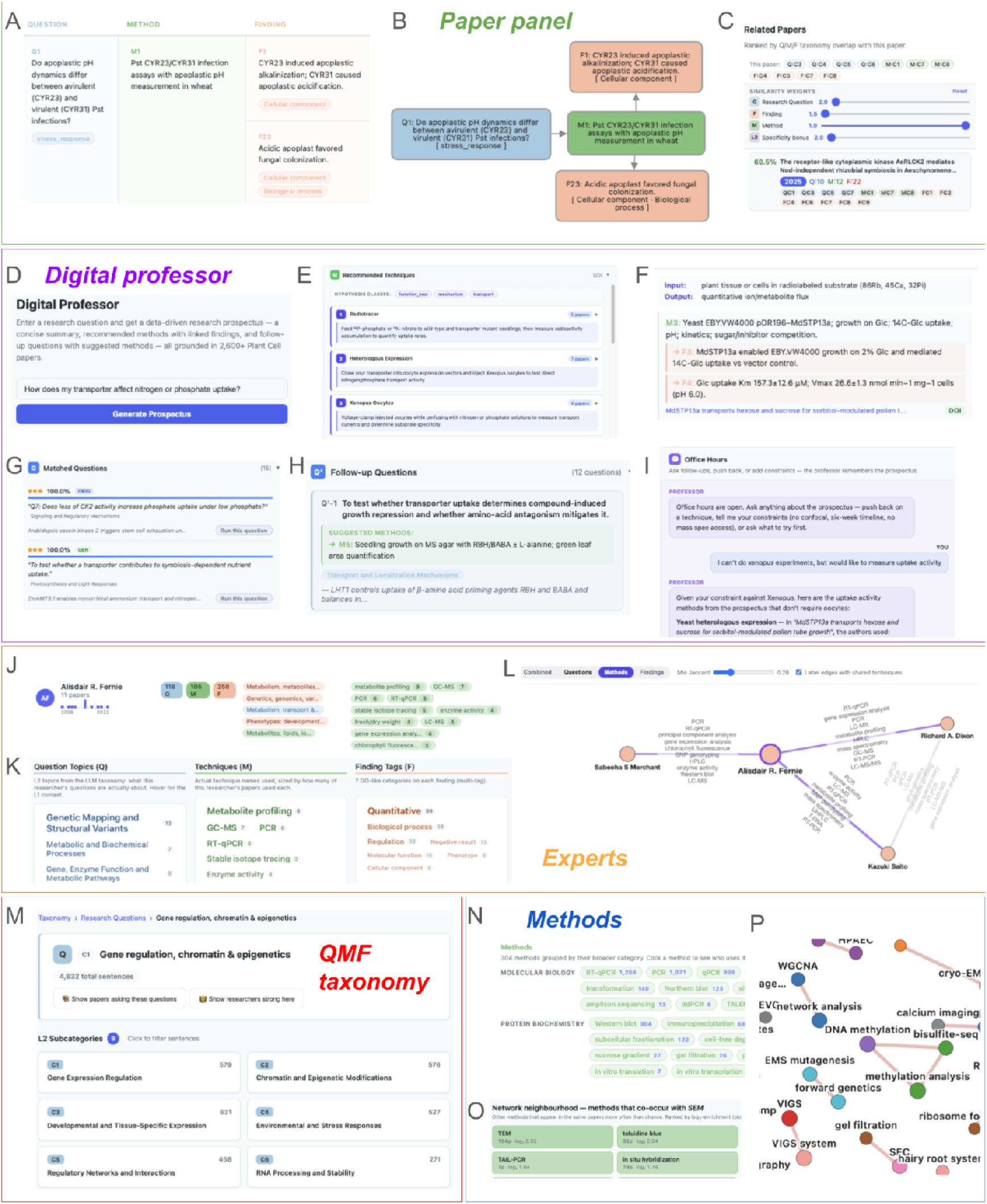
Public RPG database for browsing and reusing the Plant Cell corpus. (A–C) Paper panel. (A) Tabular Q/M/F view of a single paper. (B) Q→M→F graph view of the same paper. (C) Related-paper recommender, ranked by shared L2 Q/M/F tags; sliders allow user re-weighting of Q, F, M and L2 specificity. (D–I) Digital Professor. (D) Free-text question input that generates a research prospectus. (E) Ranked list of recommended methods with hypothesis class tags. (F) Per-method detail with input/output specification, real example M/F nodes and source DOI. (G) Corpus questions matched to the user’s query. (H) Suggested follow-up questions, each paired with its suggested method. (I) Conversational refinement of the prospectus under user constraints. (J–L) Experts. (J) Per-PI summary card (papers, Q/M/F counts, top L1 topics, top techniques). (K) Three-column speciality breakdown (Q-L2 topics, techniques, finding tags). (L) Methodological-similarity network connecting the PI to peers with overlapping technique signatures. (M) QMF taxonomy. Hierarchical browser of L1 → L2 categories with per-L2 sentence counts and links to matching papers and researchers. (N–P) Methods. (N) Method catalogue grouped by category. (O) Co-occurrence neighbourhood of a selected technique, ranked by log_2_ enrichment. (P) Force-directed similarity graph of techniques sharing paper context with the selected one.

#### Digital Professor

The Digital Professor lets users enter a natural-language research question, for example, “How does my transporter affect nitrogen or phosphate uptake?”, and returns a data-driven research prospectus grounded in our paper corpus (Figure 5D). The prospectus begins with recommended methods, ranked by the number of supporting papers (Figure 5E). Each recommendation is anchored to concrete examples drawn from real papers, including their inputs and outputs, the method nodes used and the supporting findings, with DOI links back to the source articles (Figure 5F). Alongside the user’s question, the system displays the most similar questions already asked in the corpus, ranked by similarity, here recovering a complete match to a question on CK2 and phosphate uptake from the original literature (Figure 5G). It also proposes follow-up questions with suggested methods, allowing users to plan a multi-step study rather than a single experiment (Figure 5H). Finally, an office-hours mode lets users iterate conversationally, imposing practical constraints such as the absence of confocal microscopy or Xenopus oocytes, to which the assistant adapts its recommendations (Figure 5I).

#### Experts

For any author in the corpus, an expert card summarises the author’s career span, paper count, and the numbers of Q, M, and F nodes the author contributed, together with their leading L1 question topics (Figure 5J). A three-column speciality breakdown then lists the principal investigator’s most frequent L2 question topics, the techniques they actually run (sized by number of papers) and the finding tags they accumulate (Figure 5K). A methodological-similarity network places each PI next to peers with a comparable technique signature, with edges labelled by the shared techniques, for example linking Ally Fernie and Kazuki Saito through metabolite profiling, GC-MS and PCR (Figure 5L).

#### QMF taxonomy

The full Q/M/F taxonomy is navigable as a hierarchical tree. Selecting an L1 category, such as Gene regulation, chromatin and epigenetics, exposes its L2 sub-categories with the number of generalised sentences in each and offers two pivots, “Show papers asking these questions” and “Show researchers strong here”, linking the taxonomy directly back to the paper and expert views (Figure 5M).

#### Methods

The method explorer groups all 304 techniques observed in the corpus by method category, with paper counts for each (Figure 5N). Selecting a technique, here scanning electron microscopy, returns its network neighbourhood of co-occurring methods, ranked by log2 enrichment with paper counts (Figure 5O), and renders the same neighbourhood as a technique-similarity graph so users can move from one technique to those typically used alongside it, such as from SEM to transmission electron microscopy and *in situ* hybridisation (Figure 5P).

Together, these interfaces turn the static atlas into a queryable resource, letting any researcher see in seconds what has been asked about their gene, process or organism, who works on it, and which methods the field uses to address it.

## Discussion

We have shown that the experimental logic of a research paper, the directed chain linking the Research Question asked, the Method chosen to address it, and the Finding it produced, can be extracted at scale and with high precision using large language models. Processing 2,633 Plant Cell research articles spanning two decades, we recovered typed Q, M and F nodes and assembled them into directed Q→M→F chains at greater than 98% precision against a manually annotated gold standard (Figure 1, B to D). Unlike entity-and-relation knowledge graphs, which record which biological objects are mentioned together, the RPG captures why each experiment was conducted and what it established, turning a paper’s unstructured reasoning into a machine-readable object. To our knowledge, this is the first structured map of research logic for any scientific discipline.

Beyond cataloguing individual papers, the RPG atlas allows the field itself to be studied quantitatively. We find that plant biology asks a structured rather than arbitrary set of questions, that questions are tightly coupled to particular methods and findings, and that the enriched and depleted cells of these couplings define the implicit experimental boundaries of the field (Figure 2, G and H). At the level of whole papers, this structure collapses into only seven recurring research recipes (Figure 3, A to E), within which individual investigators range from pure specialists to broad generalists (Figure 3F). Perhaps most strikingly, the strongest correlate of per-paper impact is not productivity or topical breadth but the number of distinct techniques an investigator (or their collaborative network) commands (Figure 3G), suggesting that methodological versatility, more than output volume, accompanies influential work. We emphasise that this relationship is correlational and measured across 218 investigators, and that the technique count may partly reflect laboratory size, resources, and the collaborative network; it should therefore be read as a hypothesis about how impactful research is conducted rather than as a prescription. The atlas also revises a common intuition about methodological progress: across twenty years, plant biology has turned over its peripheral techniques, replacing PCR, Northern blot and microarray with RNA-seq, CRISPR and ChIP-seq, while a stable core of methods persisted and the number of techniques and method categories per paper barely increased (Figure 4, A to D). Plant biology has therefore re-tooled its bench without broadening it.

As with any LLM-based extraction, the RPG atlas is accurate but not infallible, and it inherits limitations that warrant attention. The pipeline can occasionally misunderstand a node or attach an edge between unrelated QMF nodes. These errors could be reduced by fine-tuning the models on curated examples of the expected output, and users are encouraged to follow the DOI links provided in every interface to verify a node or chain against the source article. A more fundamental limitation is one of scope. Our atlas rests on a single journal, thus reflects Plant Cell’s editorial focus, model-organism emphasis and methodological culture. The recipes, couplings and trends reported here should accordingly be read as properties of this corpus rather than of plant science as a whole. The pipeline is also currently restricted to the information contained in the analysed text, so questions, methods, and findings reported only in figures, supplementary materials, or full-text sections beyond our input may be underrepresented.

The central strength of the RPG framework, however, is that it is journal- and field-agnostic, and our future efforts will focus on testing it across the entire plant-science literature. By extending the pipeline to all plant science papers, drawing on other plant journals, and by incorporating full-text articles rather than abstracts alone, we aim to build a comprehensive, cross-journal atlas of plant biology. Such a resource would let us ask whether research recipes and question-method-finding couplings are conserved across the field or are specific to individual communities, track how questions and techniques propagate between sub-fields over time, and surface understudied questions and underused methods that no single journal makes visible. As the corpus grows, the accompanying public database and its literature-grounded Digital Professor will become an increasingly complete map of what plant biology has asked, how it has answered, and where its open questions remain. We anticipate that the RPG atlas will become a valuable resource for the plant science community, supporting activities ranging from orienting a newcomer to an unfamiliar topic to planning the next experiment in an established research programme.

## Materials and methods

### Procurement of full-text research articles

Full-text Plant Cell articles (2005–2026) were scraped as html files organised by year and month, and each file’s header was parsed to extract its DOI, title, and source URL (n = 3,366 articles). To restrict the corpus to primary research, every DOI was resolved to a PubMed PMID via NCBI E-utilities esearch, and the corresponding MEDLINE record was retrieved with efetch. Articles whose publication-type (PT) lines included Review, Editorial, Comment, Letter, News, Biography, Case Reports, Historical Article, Congresses, or Consensus Development Conference were excluded, retaining 2,301 experimental papers (68.4% of the scraped set; 316 articles without a resolvable PMID were dropped). The corpus was subsequently expanded with more recent issues to yield the final 2,633 experimental Plant Cell papers used in this study. Of these, 2,499 have a last-author PI identified (94.9%) and 2,583 have a citation count from OpenAlex (98.1%)

### Construction of the RPG database

The RPG was built from the full text of every research article in *The Plant Cell* via a four-stage LLM pipeline: (1) per-paper *Q*→ *M*→ *F* graph extraction, (2) generalisation of paper-specific labels into corpus-comparable statements, (3) bottom-up L1 taxonomy induction, and (4) per-L1 L2 sub-taxonomy induction.

### Per-paper *Q*→*M*→ *F* graph extraction

For each paper, the cleaned experimental text was submitted to the Responses endpoint of GPT-5 (reasoning effort = “high”). A single prompt instructed the model to return a JSON object containing the paper title, a list of nodes labelled *Q* (Research Question), *M* (Method), or *F* (Finding), and a list of directed edges restricted to *Q*→ *M* and *M*→ *F*. Node labels were required to be near-verbatim, ≤ 25 words, and uniquely identified (Q1, Q2, …, M1, M2, …, F1, F2, …). Prompt-level instructions enforced edge-building rules that favour aggregation: (i) any single method shared across multiple questions reuses one *M*-node rather than spawning duplicates, producing many-to-one *Q* → *M* fan-in; (ii) one method that yields multiple findings produces one-to-many *M* → *F* fan-out; (iii) the same assay, software pipeline, or sequential wet-lab protocol block is grouped into a single *M*-node; (iv) every *F*-node must point back to at least one *M*-node and every *M*-node must originate from at least one *Q*-node; (v) the final graph must be acyclic. Negative results were retained as valid *F*-nodes. Outputs were validated as JSON and saved one file per paper.

### Generalisation of node labels

To enable cross-paper comparison, each per-paper node JSON was passed through a second GPT-5-mini call (reasoning effort = “high”) together with that paper’s abbreviation glossary. The prompt converted the paper-specific verbatim label of each *Q, M*, or *F* into a *generalised form*: a stand-alone scientific statement intelligible without the source paper’s gene names, sample identifiers, or local abbreviations. The output of this stage is a parallel JSON tree containing, for every node, both the original label and its corpus-comparable generalisation.

### L1 taxonomy induction

A top-level taxonomy was induced bottom-up for each node type (*Q, M, F*) independently. All generalised statements of a given type were collected across the entire corpus, randomly shuffled (seed = 42 for reproducibility), and partitioned into batches of 120. Each batch was submitted to GPT-5.2 (temperature = 0.2, JSON-mode) with instructions to propose 6–12 top-level categories and assign every input sentence to exactly one category id. The resulting per-batch taxonomies were merged hierarchically: batches were grouped into “super” chunks of 10, each super-merge produced 6–15 categories, and a final super-to-final LLM call yielded the canonical L1 taxonomy. At every merge level the model was required to return a merged_to_inputs mapping such that each input category appears in exactly one output category, which is a constraint we verified programmatically against the universe of all batch-local category ids. Categories missed by the merge were collected into recovery buckets (C_RECOVERED and C_RECOVERED_BATCH) and re-assigned to an existing final category in a separate GPT-4o pass (temperature = 0.0); any final mapping with zero missing entries passed the audit. The output of this stage is, per node type, a final taxonomy ({*categories*: [{*id, name, description*}]}), a final-to-batch index, and an enriched table that joins each final category to all its supporting generalised sentences along with paper-level provenance.

### L2 taxonomy induction

For each pair (node type, L1 category) the procedure above was repeated at finer granularity. Generalised sentences assigned to the parent L1 were partitioned into batches of 200 and classified with GPT-4o (temperature = 0.2) into exactly 10 L2 sub-categories per batch. Per-batch taxonomies were merged hierarchically (chunk size = 10, target 6–12 final categories) using GPT-4o at temperature 0.0. The same merged_to_inputs completeness constraint, batch-level coverage validation, and recovered-category patch step were applied at L2. Batch classification calls were parallelised across 50 concurrent workers within each L1 category. The procedure yields, per (type, L1) pair, a final L2 taxonomy with sentence-level provenance and a per-category sentence count. Table S4 lists the model, temperature, batch unit, and concurrency mode of every pipeline stage.

### Similarity measures

The RPG database employs four distinct similarity or association measures, each tuned to the granularity of the entities being compared.

### Paper-to-paper similarity

Each paper *p* is represented as a categorical fingerprint formed by aggregating its *Q, M*, and *F* nodes into six sets,

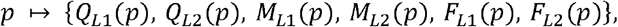

where *X*_*Li*_ (*p*) is the set of L1 (or L2) taxonomy categories assigned to nodes of type *X* ∈ {*Q, M, F*} in paper *p*. Pairwise similarity between papers *a* and *b* is a weighted average of dimension-wise Jaccard coefficients,

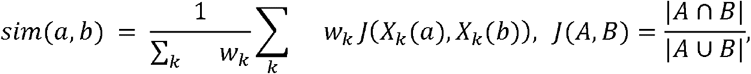

with *k* indexing the six dimensions. Default weights are *w*_*Q,L*1_ = 2.0, *w*_*F,L*1_ = 1.5, *w*_*M,L*1_ = 1.0, all multiplied by an L2-specificity bonus *ℓ*= 2.0 to obtain the L2 weights (*w*_*X,L*2_ = *w*_*X,L*1_ *ℓ*). The four parameters are exposed as user-tunable sliders.

### Researcher-to-researcher similarity

Each researcher *r* is represented by three sparse non-negative count vectors over the taxonomy,

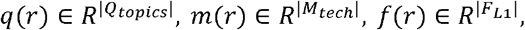

where *q*_*i*_ is the contribution count of researcher *r* to Q-topic *i* (sum across the researcher’s papers of *Q*-nodes mapped to that L2 topic), *m*_*i*_ is the number of the researcher’s papers tagged with technique *i*, and *f*_*i*_ is the contribution count to F L1 category *i*. Pairwise similarity for each modality is the weighted Jaccard (Ruzicka) index,

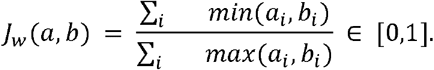

The combined score is the arithmetic mean of the three modal Jaccards,

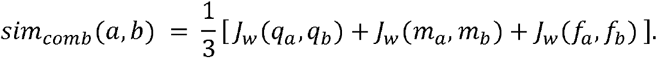

### Method co-occurrence enrichment

For each unordered pair of techniques (*a, b*), observed and expected co-occurrence counts are

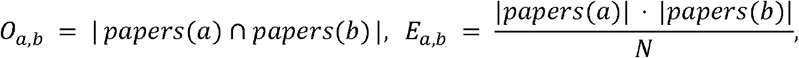

where *N* is the size of the union of all papers carrying any method tag. The pairwise enrichment is the base-2 log-ratio with Laplace smoothing (*α* = 0.5) to bound variance on rare pairs,

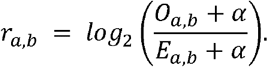

Edges with *O*_*a,b*_ < 2 are discarded as noise. Positive *r*_*a,b*_ indicates over-representation (methods co-occur more often than expected by chance); negative values indicate avoidance. The same statistic powers both the global method co-occurrence network on the Methods tab and the per-method “Network neighbourhood” panel on each method’s detail page.

### Implementation

All similarity structures are pre-computed once at server start-up and held in memory. The researcher-similarity tensor across *N* ≈ 1,500 authors and ∼ 305 techniques is constructed in under five seconds on a single core using inverted-index lookups over Python dict/set primitives. The method co-occurrence network is recomputed live on request from the technique-to-papers index. All endpoint paths and source files are documented in the project repository.

## Data availability

RPG database source code and scripts to generate the RPG is available at: https://github.com/manojitharaju016/rpg_project. The Obsidian wiki descripting the RPG concepts are available at: https://publish.obsidian.md/mutwillab/Homepage/Publication+Vaults/RPG/README

## Acknowledgments

The authors wish to thank the members of the Mutwil lab for their support, encouragement and comments on the manuscript.

## Author Contributions

Jing Yang led the RPG construction, evaluation, and taxonomy identification. Manoj Itharajula led web development, cloud deployment, user interface enhancement and co-supervised Jing Yang. The project was conceptualised and supervised by Marek Mutwil.

## Supplemental Data

### Supplemental Tables

**Table S1. Each row corresponds to one paper from The Plant Cell included in the Research Process Graph (RPG)**. Columns are organised in four blocks: (i) Paper metadata: paper_id (internal 12-character hash), title, year, month, doi, corresponding_authors (semicolon-separated “Given Family” list, last authors as reported by Crossref), and citation_count (Crossref is-referenced-by-count at the time of retrieval; blank if unavailable, not_in_crossref if the DOI returned HTTP 404). (ii) L1 taxonomy assignment (32 columns): one binary indicator per top-level (L1) category of the Question (Q), Method (M), and Finding (F) taxonomies. Column names follow <X>_L1 <categoryID>__<slug>, where X ∈ {Q, M, F}. A cell is 1 if at least one Q/M/F node of paper p was assigned to that L1 category by the asymmetric token-coverage classifier (top-2 L1 per node, threshold cov > 0), and 0 otherwise. (iii) L2 taxonomy assignment (271 columns): one binary indicator per L2 sub-category, with column names <X>_L2 <parentL1> <slug>. Encoded analogously to (ii) using top-2 L2 assignments per node; any L2 hit additionally propagates its parent L1 into block (ii). (iv) Technique fingerprint (305 columns): one binary indicator per generalised laboratory/computational technique (tech <slug>), derived from the M-node technique index (m_node_techniques_v2.json). A cell is 1 if the paper contains at least one M-node tagged with that technique. Totals: 7 metadata + 32 L1 + 271 L2 + 305 technique = 615 columns × 2,633 rows. The full algorithmic definition of node→taxonomy assignment and the four-stage construction pipeline are described in the Materials and Methods.

**Table S2. Node-level evaluation of LLM-extracted Q/M/F nodes from a representative artificial intelligence paper**. Each row corresponds to a single node extracted by one of three LLMs (GPT-5, GPT-5-mini, GPT-OSS-120B). Columns report the node type identifier (Q/M/F with index), the semantic subtype (e.g., research question, experimental aim, core method, specific result), the extracted text, a correctness score (1 = correct, 0.5 = uncertain, 0 = incorrect), an explicitness score (1 = explicitly stated in the paper, 0.5 = ambiguous, 0 = implicit or inferred by the model), the LLM that produced the node, and optional evaluator comments. Correctness was assessed by comparing each extracted node against the original paper text: a node was scored as correct if its content could be directly traced to a statement in the paper, incorrect if it contained factual errors or hallucinated content, and uncertain if it had a plausible semantic association with the paper content but could not be unambiguously confirmed or rejected. The winner column indicates the best-performing model for this paper.

**Table S3. Node-level evaluation for a representative computational biology paper**. Format and scoring criteria as in Table S2.

**Table S4. Node-level evaluation for a representative cell biology paper**. Format and scoring criteria as in Table S2.

**Table S5. Node-level evaluation for a representative genomics paper**. Format and scoring criteria as in Table S2.

**Table S6. Node-level evaluation for a representative metabolism and physiology paper**. Format and scoring criteria as in Table S2.

**Table S7. Edge-level evaluation of Q**→**M**→**F chains from a representative artificial intelligence paper**. Each row corresponds to a single Q→M→F chain extracted by one of two LLMs (GPT-5, GPT-5-mini). Columns report the chain number, the full text of the Q, M, and F nodes forming the chain, and three relationship scores: Q→M (1 = correct, 0.5 = uncertain, 0 = incorrect), M→F (same scale), and Q→M→F (same scale), along with the LLM name and optional evaluator comments. The Q→M relationship was scored as correct if the method was logically applied to address the stated question in the original paper; the M→F relationship was scored as correct if the finding was a direct or clearly supported outcome of the stated method. A complete Q→M→F chain was scored as correct only when both the Q→M and M→F relationships were independently correct; if either was uncertain or incorrect, the full chain received the lower score. Edge evaluation results are reported as proportions (fraction of correct, uncertain, and incorrect edges per model) rather than raw counts, because one-to-many connections in the DAG structure cause the number of chains to vary substantially between papers, and proportions enable fairer cross-model comparison.

**Table S8. Edge-level evaluation for a representative computational biology paper**. Format and scoring criteria as in Table S7.

**Table S9. Edge-level evaluation for a representative cell biology paper**. Format and scoring criteria as in Table S7.

**Table S10. Edge-level evaluation for a representative genomics paper**. Format and scoring criteria as in Table S7.

**Table S11. Edge-level evaluation for a representative metabolism and physiology paper**. Format and scoring criteria as in Table S7.

**Table S12. One row per principal investigator (last-author) of at least one corpus paper**. Researcher identity and paper assignment are inherited from the Crossref-derived last-author cache and the corresponding OpenAlex records. The seven research recipes (R1–R7) are the k-means clusters defined in Figure 3A; the dominant recipe and recipe entropy summarise how each PI’s papers distribute across them. Q/M/F L1 and L2 categories follow the hierarchical taxonomy in Figure 2; cells list the top categories together with their node counts in the format “Category (count)”, with multiple entries separated by semicolons. Columns are: researcher_id, internal identifier; name, family, given, orcid, author identity; paper_count, number of corpus papers for which the researcher is last-author; year_first, year_last, span of those papers; n_Q_nodes, n_M_nodes, n_F_nodes, total Question, Method and Finding nodes contributed; recipe_entropy, Shannon entropy (log_2_) of the PI’s recipe distribution, ranging from 0 (pure specialist) to log_2_(7) ≈ 2.81 (full generalist); dominant_recipe, most frequent recipe (R1–R7) with its Q-L1 → M-L1 → F-L1 triad; recipe_distribution, paper counts per recipe, sorted from most to least frequent; Q_L1, top 5 Question L1 categories; Q_L2, top 8 Question L2 categories shown as “L1 > L2”; M_L1, M_L2, F_L1, F_L2, the equivalent breakdowns for Method and Finding nodes; method_categories, top 5 high-level method categories from the curated technique dictionary; techniques, top 10 specific techniques (e.g. RT-qPCR, Western blot, confocal microscopy) the PI employs across their corpus papers.

**Table S13. Models, temperatures, and concurrency modes used by the RPG construction pipeline**.

### Supplemental Figures

**Figure S1.**
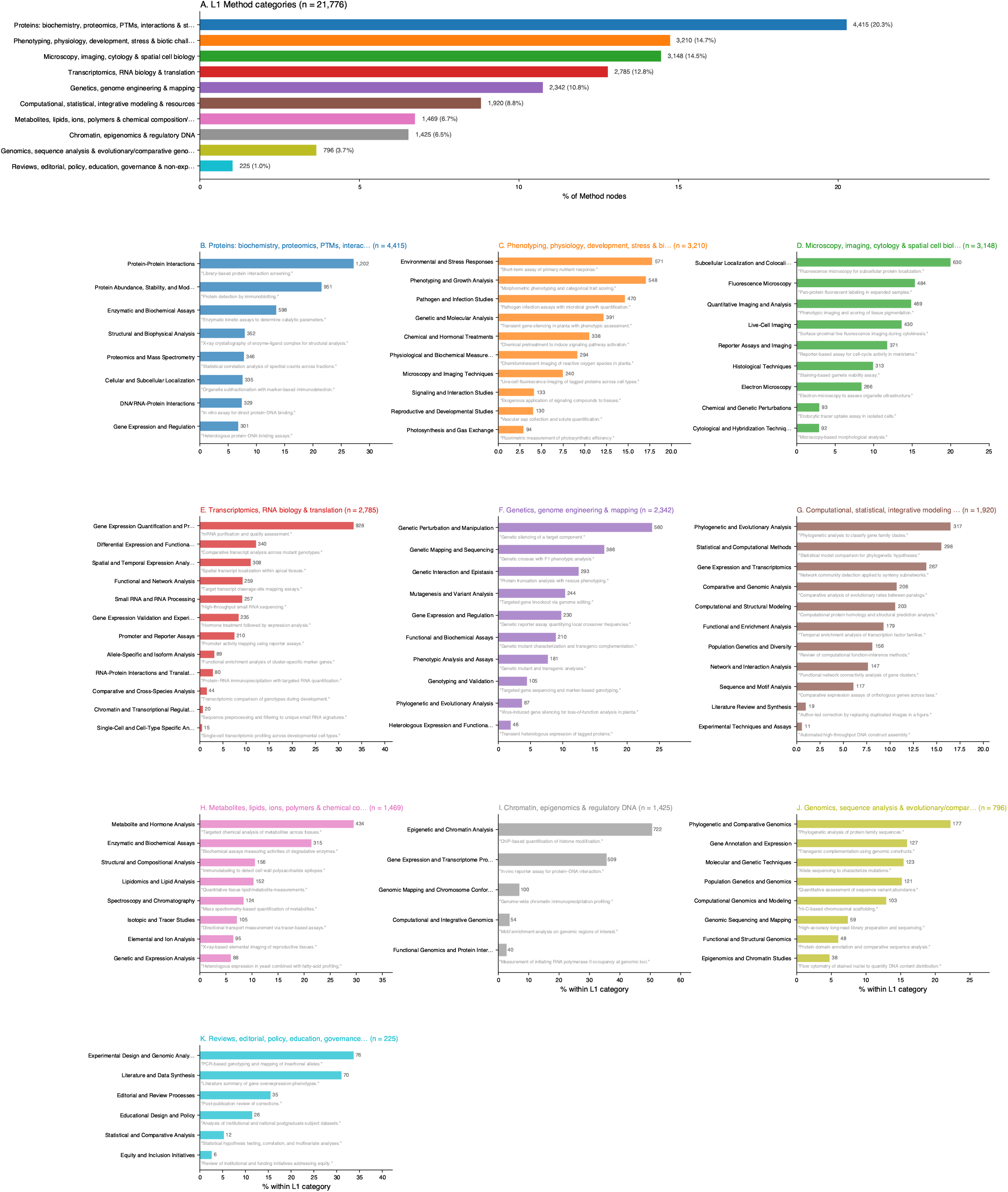
Full L1 + L2 taxonomy of Question nodes across the Plant Cell corpus. (A) L1 overview: number of Question (Q) nodes assigned to each of the 10 L1 categories (n = 23,456 Q nodes after removing “Recovered Batch/Miscellaneous”), with absolute counts and percentages annotated to the right of each bar. (B–K) Per-L1 zoom panels showing the L2 sub-category breakdown of every L1 category, ordered by L1 size. Within each panel, bars give the percentage of Q nodes within that L1 falling into each L2 sub-category; the absolute node count is shown next to each bar. A single representative generalised question is printed in grey italics below every L2 bar to illustrate the kind of sentence that the L2 category collects. Panel colours match the corresponding L1 bar in (A).

**Figure S2.**
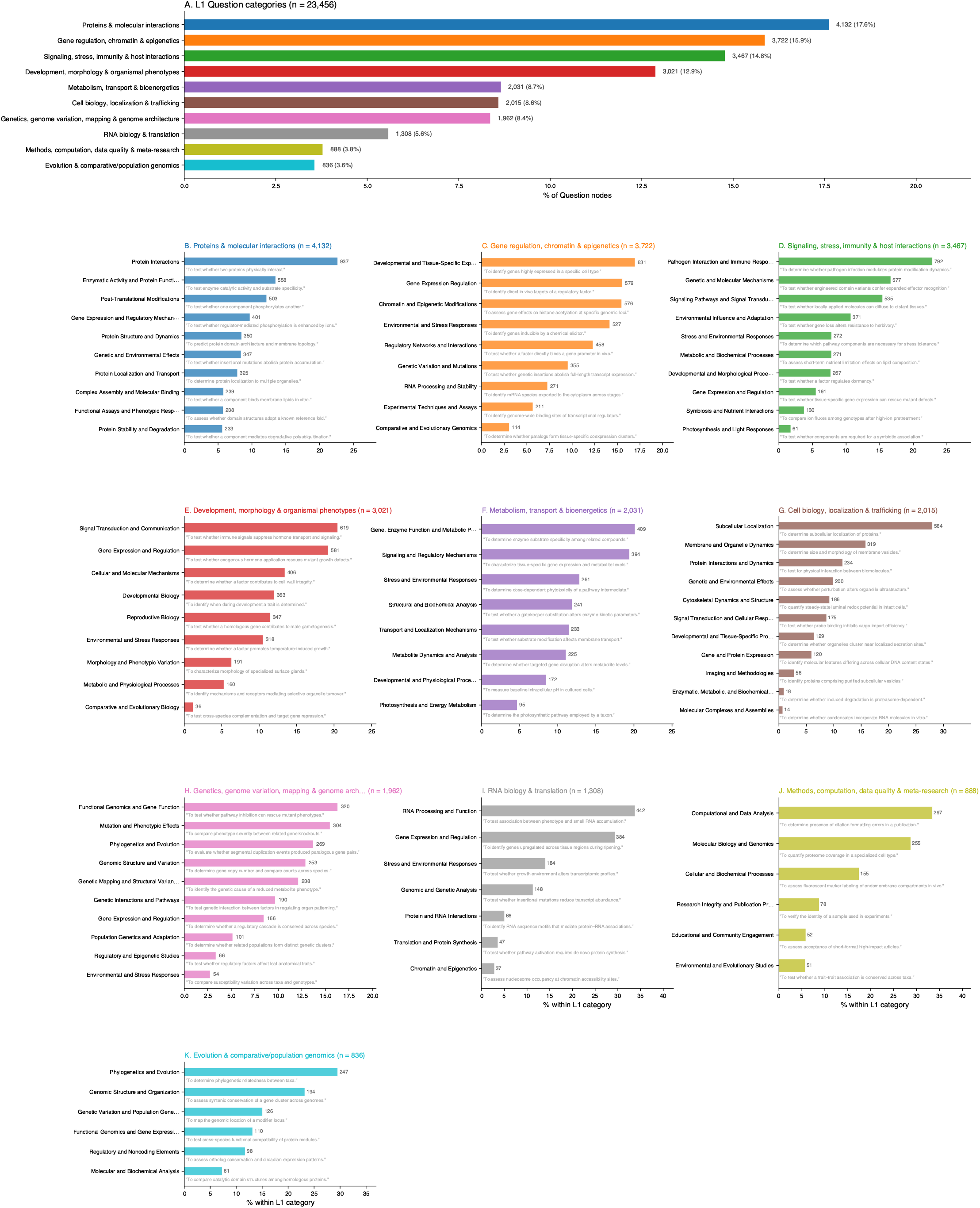
Full L1 + L2 taxonomy of Method nodes across the Plant Cell corpus. (A) L1 overview: number of Method (M) nodes assigned to each of the 10 L1 categories (n = 21,776 M nodes after removing “Recovered Batch/Miscellaneous”), with absolute counts and percentages annotated to the right of each bar. (B–K) Per-L1 zoom panels showing the L2 sub-category breakdown of every L1 category.

**Figure S3.**
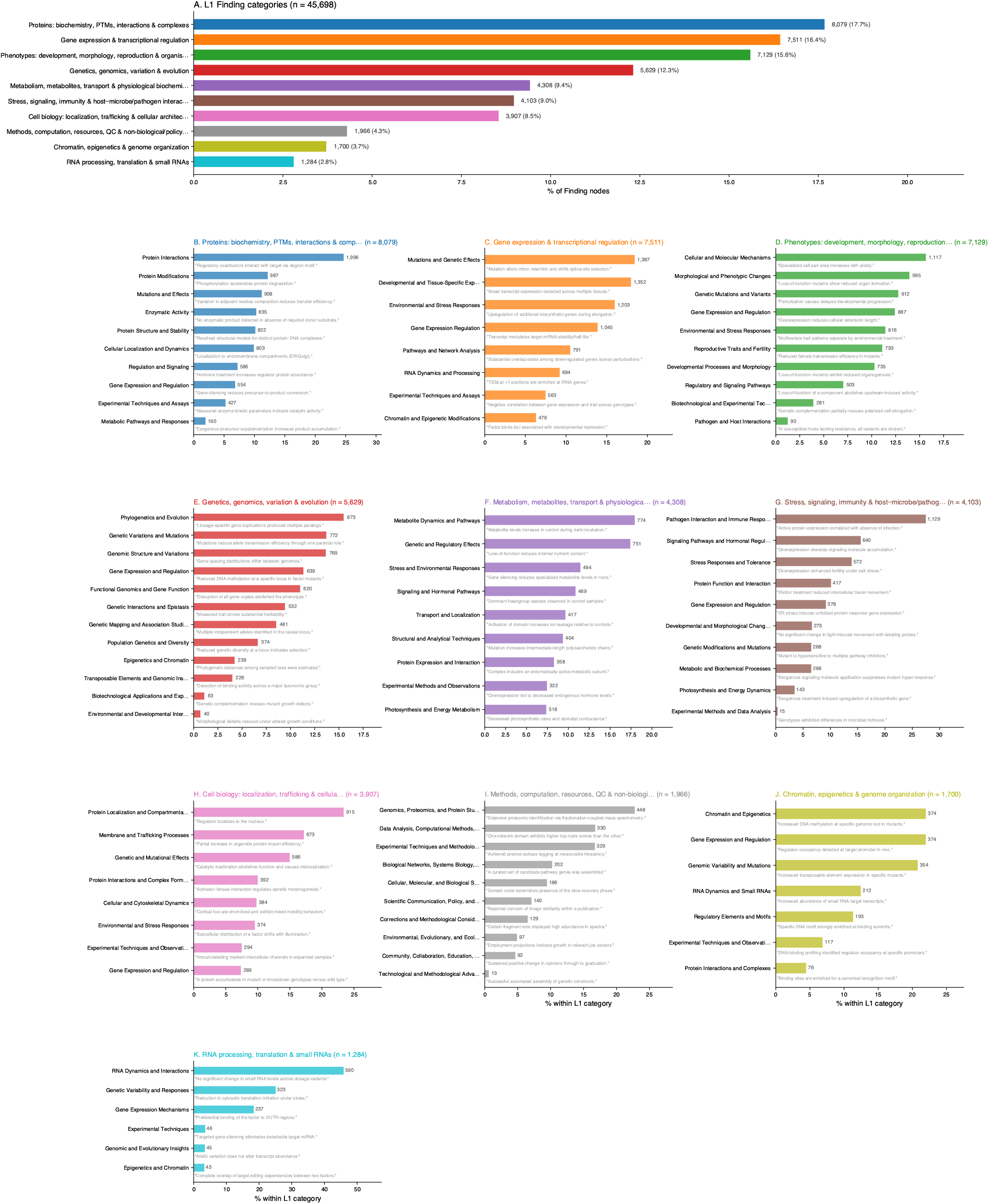
Full L1 + L2 taxonomy of Finding nodes across the Plant Cell corpus. (A) L1 overview: number of Finding (F) nodes assigned to each of the 10 L1 categories (n = 45,698 F nodes after removing “Recovered Batch/Miscellaneous”), with absolute counts and percentages annotated to the right of each bar. B–K) Per-L1 zoom panels showing the L2 sub-category breakdown of every L1 category.

**Figure S4.**
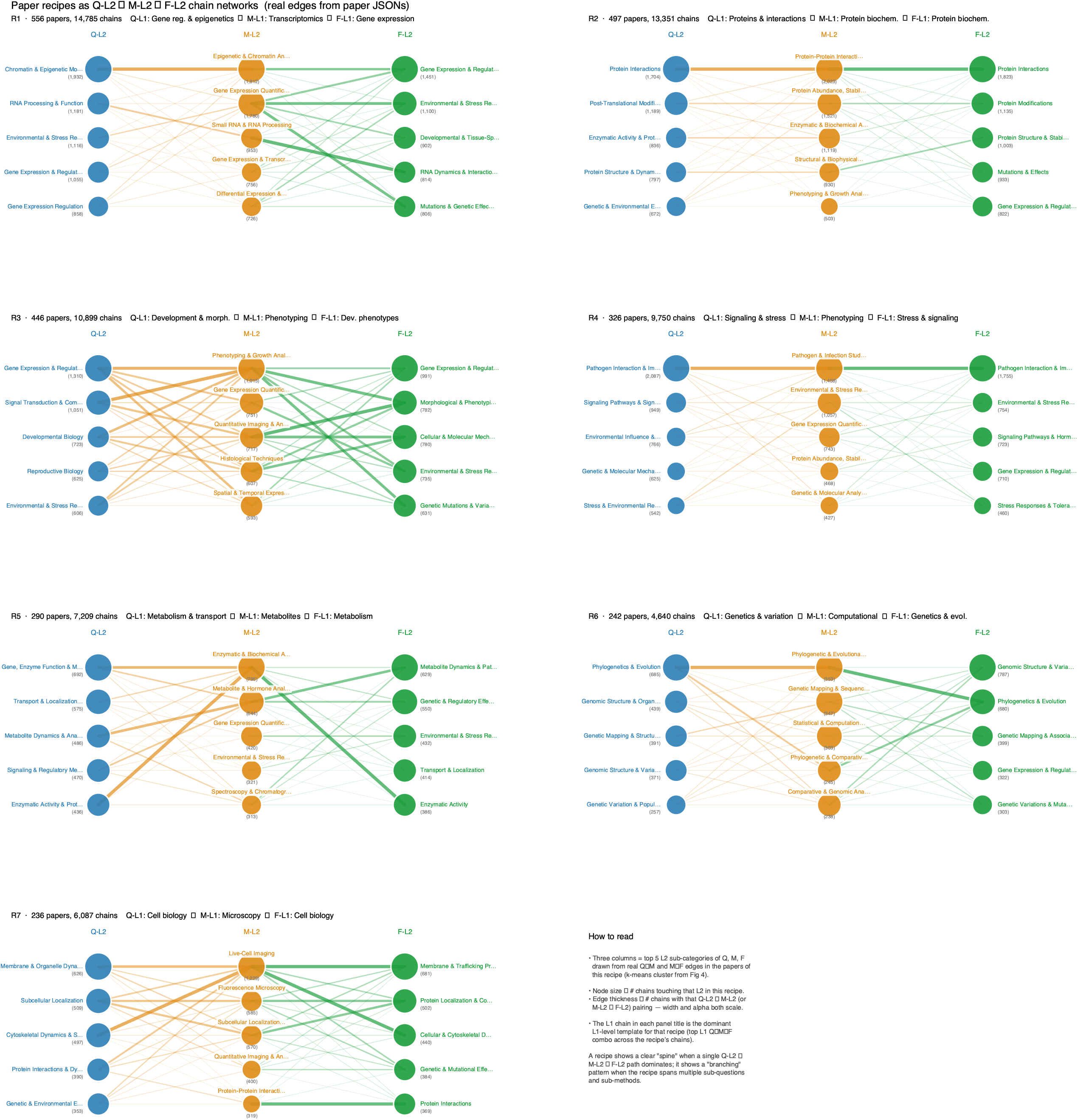
Recipes for the seven major paper categories.

